# Solutions of ferrous salts protect liposomes from UV damage: implications for Life Origin

**DOI:** 10.1101/2024.04.29.591450

**Authors:** Ben Turner, Gennady Fiksel, Vladimir Subbotin

## Abstract

Previously, we presented a hypothesis on the Darwinian evolution of liposomes that relies only on natural and ever-present phenomena: solar UV radiation, day/night cycle, gravity, and the formation of amphiphiles, e.g., fatty acids and phospholipids, in aqueous media. The hypothesis further suggests that liposomes, formed at the air-water interface, will be inevitably destroyed by Sun UV unless they are submerged and shielded by primordial water. The hypothesis makes two testable predictions. First, certain ingredients of the Archean waters, e.g., ferric salts, can completely attenuate UV; and second, such ferric salts solution can protect the submerged liposomes from the solar UV damage. In our previous experiments (Subbotin and Fiksel 2023a), we tested the prediction of UV attenuation by ferrous salt solutions. We have demonstrated that out of several tested iron salt solutions, two salts demonstrated very strong UV attenuation: at a concentration of 2.5 g/L of (NH_4_)_5_[Fe(C_6_H_4_O_7_)_2_] and FeCl_3_, the UV intensity drops by a factor of 1000 at a submersion depth of 7.4 mm and 6.3 mm, respectively. In the present study, we tested the second prediction: we investigated whether the above ferrous salt solutions can protect UV-sensitive liposomes from UV damage. The results demonstrated that 10 mm of solutions, the ferric ammonium citrate or iron trichoride, completely protected UV-sensitive liposomes from UV damage. These results supplement and reinforce the proposed hypothesis.

## Introduction: Solar UV and the origin of life

“One of the many paradoxes encountered in the early history of life lies in the fact that the same rays of the Sun which formed the building blocks of the molecules of life were lethal for life. Early life had, therefore, only limited environmental possibilities. It could be survived only when shielded by a thick layer of water…”, Martin Rutten, ‘Origin of life by natural causes’(Rutten 1971).

In our previous communication, we considered the Archean water-air interface as a spatial plane crucial for the emergence of abiogenesis, namely liposomes, which at the same time appear as an obvious subject of destruction by solar UV. Following the idea of Martin Rutten, we suggested that the liposomes encapsulate a solute heavier than the surrounding media will descend from the water-air interface and be protected from UV by a layer of water. We also calculated the time and depth of the submergence, both of which depend on known physical characteristics: liposomal size and specific gravity (Subbotin and Fiksel 2023b). However, the UV-shielding ability of the presumable Archean water depends on its composition and remains a subject of experimental tests.

In the follow-up experiments(Subbotin and Fiksel 2023a), we examined the UV-shielding ability of several ferrous salts: salts (FeCl_2_—iron dichloride, FeCl_3_—iron trichoride, Fe(NO_3_)_3_—ferric nitride, NH_4_Fe(SO_4_)_2_—ferric ammonium sulfate, and (NH_4_)_5_[Fe(C_6_H_4_O7)_2_]—ferric ammonium citrate), which were suggested to be present in the Archean waters (Breslow and others 2010; Gómez and others 2007; Handschuh and Orgel 1973; Hardy and others 2015; Keefe and Miller 1996; Kim and others 2016; Miller and Urey 1959; Patel and others 2015; Sproul 2015). All salts were dissolved in water at 2.5 g/L, a concentration previously suggested for the Archean waters (Gómez and others 2007). The results showed that solutions of two salts, (NH_4_)_5_[Fe(C_6_H_4_O_7_)_2_] and FeCl_3_, demonstrated very strong UV attenuation: the UV intensity drops by a factor of 1000 at a submersion depth of 7.4 mm and 6.3 mm, respectively.

In the present study, we investigated whether specially designed liposomes with membranes containing UV-sensitive polymers (Ahn and others 2009) could be affected by UV, and, if so, whether the solutions of (NH_4_)_5_[Fe(C_6_H_4_O_7_)_2_] and FeCl_3_ can reduce the UV damaging effect and protect liposomes.

## Materials and Methods

### Liposomes

We used large polymerizable UV-sensitive multilamellar liposomes composed of DBPC:PCDA (80:20) molar ratio with a total lipid concentration of 7 mM in PBS buffer at pH 7.4 (Encapsula NanoSciences, Brentwood, TN, USA). In all experiments, 100 μl of undiluted liposome suspension was added to wells. Upon UV radiation of the liposomes, the membrane undergoes destruction, and colorimetrically sensitive membrane polymers change color from white to blue (Ahn and others 2009).

#### UV source and radiation setup

We used a short-wave (254 nm) bactericide UV lamp placed 30 mm above a 96-well plate containing liposomes. The plate was covered with UV-impermeable lead to block UV radiation from undesired wells, with holes only above the designated wells. The lamp warm-up time was 1 min, and the radiation time was 3 min.

#### Setup of UV shielding device

The disposable semi-micro UV cuvettes, fully filled with (NH_4_)_5_[Fe(C_6_H_4_O_7_)_2_] or FeCl_3_ solutions, were hermetically sealed by Parafilm M sealing film. The cuvettes’ material is transparent to UV, so the UV absorption was entirely due to the composition of the solutions. The inside dimension of the upper part of the cuvette is 10x10x20 mm, which completely covers two adjacent wells. Cuvettes filled with distilled water were used as non-attenuated UV controls.

#### Assessment of the liposomal damage and protection

We used the BioTek Synergy Neo2 Hybrid Multimode Reader (Agilent, Santa Clara, CA, USA) to measure the colorimetric shift of the UV-sensitive liposomes/ polymerizable polymers. Alternatively, we recorded results by visually assessing the liposomes’ color change.

## Results

### Multimode Reader data

The results clearly demonstrated a shift in the color, observed both by the spectrometer and visually, of liposomes covered by cuvettes with distilled water after 3 min UV radiation. On the contrary, liposomes covered by cuvettes with solutions either (NH_4_)_5_[Fe(C_6_H_4_O_7_)_2_] or FeCl_3_ retained the original color – see Fig. 1 and Fig. 2.

**Figure 1.**
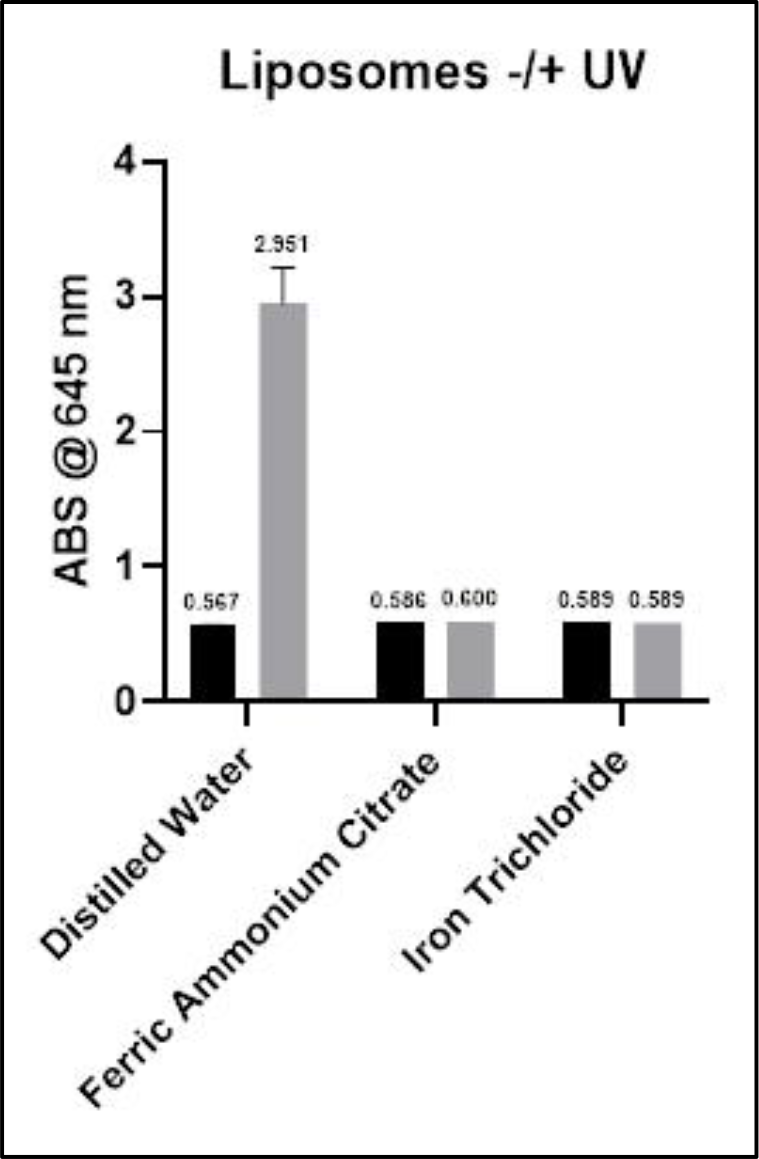
Absorbance data was acquired on a Perkin Elmer Synergy Neo2 plate reader equipped with monochromator optics. We first conducted spectral scanning from 380 nm to 740 nm (5 nm steps) to determine the wavelength of maximal OD increase (645 nm). Subsequent endpoint reads were acquired pre- and post-UV exposure at that single wavelength. Black columns – liposomes before the UV irradiation; grey columns – liposomes radiated with UV for 3 minutes.

**Figure 2.**
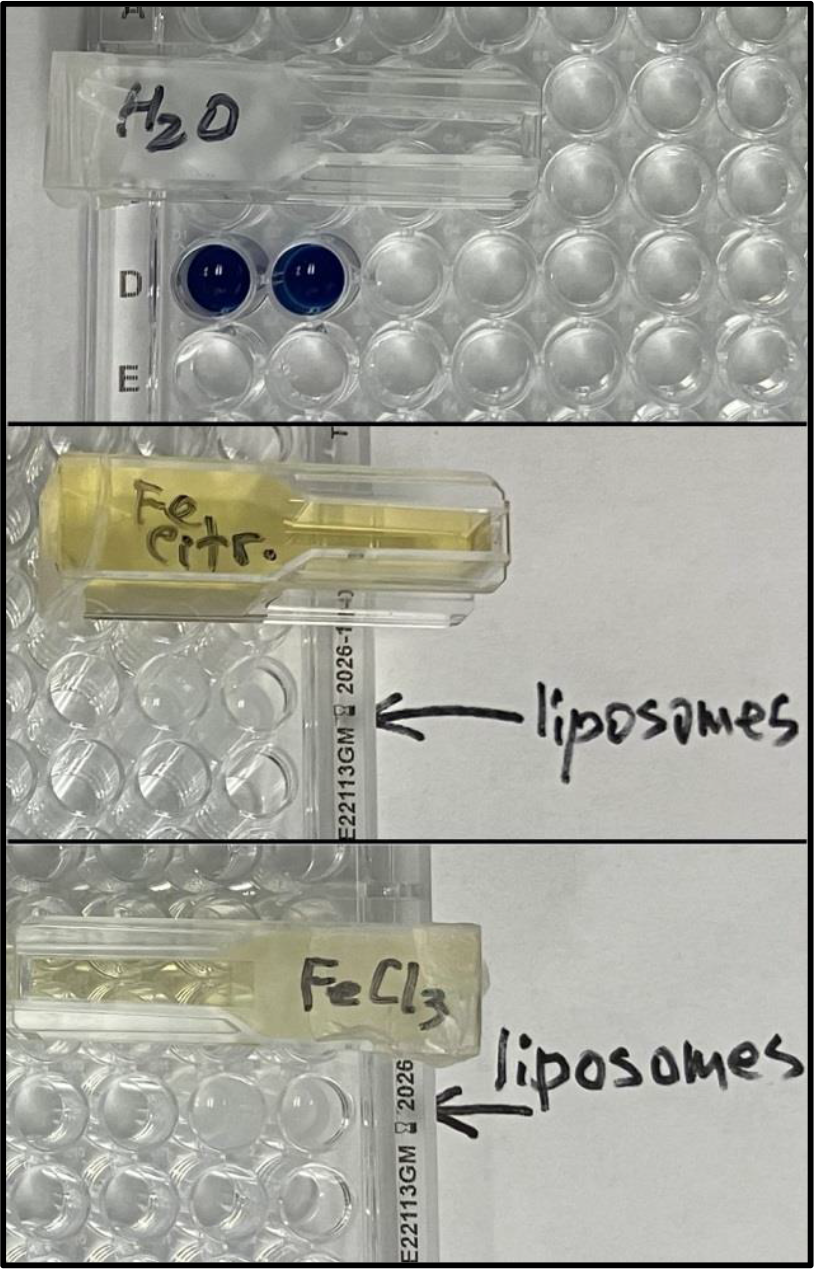
Liposomes after 3 min of UV radiation. The top image shows liposomes shielded by cuvettes with distilled water, and the middle and bottom images show liposomes shielded by cuvettes with solutions of (NH_4_)_5_[Fe(C_6_H_4_O_7_)_2_] and FeCl_3_.

Images of liposomes after UV radiation with shielding by water or ferrous salts solutions also demonstrated a dramatic shift in liposomes’ color - Fig. 2.

### Summary and conclusions

The results of this study demonstrated that: 1) Liposomes can be destroyed by UV after as short radiation time as 3 min; 2) a thin layer (10 mm) of ferrous salt solutions, (NH_4_)_5_[Fe(C_6_H_4_O_7_)_2_] and FeCl_3,_ completely protected liposomes from UV destruction.

The experimental results confirmed our suggestion that the ferric ammonium citrate and iron trichoride solutions could constitute a protective media for life origin within the Lipid World scenario (Subbotin and Fiksel 2023a, and reinforced our hypothesis (Subbotin and Fiksel 2023b) that Sun UV radiation could be one of the major selection forces at the abiogenesis stage of life origin.

## Acknowledgment

The authors want to thank Drs. Sara Mir (Encapsula NanoSciences, Brentwood, TN) and David Rozema (Empirico Inc., Madison, WI) for valuable advice and technical support.

## References

Ahn DJ, Lee S, and Kim JM. (2009) Rational design of conjugated polymer supramolecules with tunable colorimetric responses. Advanced Functional Materials, 19, 1483–1496.

Breslow R, Levine M, and Cheng Z-L. (2010) Imitating prebiotic homochirality on Earth. Origins of Life and Evolution of Biospheres, 40, 11–26.

Gómez F, Aguilera A, and Amils R. (2007) Soluble ferric iron as an effective protective agent against UV radiation: Implications for early life. Icarus, 191, 352–359.

Handschuh G, and Orgel L. (1973) Struvite and prebiotic phosphorylation. Science, 179, 483–484.

Hardy MD, Yang J, Selimkhanov J, Cole CM, Tsimring LS, and Devaraj NK. (2015) Self-reproducing catalyst drives repeated phospholipid synthesis and membrane growth. Proceedings of the National Academy of Sciences, 112, 8187–8192.

Keefe AD, and Miller SL. (1996) Was ferrocyanide a prebiotic reagent? Origins of Life and Evolution of the Biosphere, 26, 111–129.

Kim HJ, Furukawa Y, Kakegawa T, Bita A, Scorei R, and Benner SA. (2016) Evaporite borate-containing mineral ensembles make phosphate available and regiospecifically phosphorylate ribonucleosides: borate as a multifaceted problem solver in prebiotic chemistry. Angewandte Chemie, 128, 16048–16052.

Miller SL, and Urey HC. (1959) Organic compound synthesis on the primitive earth. Science, 130, 245–51.

Patel BH, Percivalle C, Ritson DJ, Duffy CD, and Sutherland JD. (2015) Common origins of RNA, protein and lipid precursors in a cyanosulfidic protometabolism. Nat Chem, 7, 301–7.

Rutten MG. (1971) Origin of life by natural causes.

Sproul G. (2015) Abiogenic syntheses of lipoamino acids and lipopeptides and their prebiotic significance. Origins of Life and Evolution of Biospheres, 45, 427–437.

Subbotin V., and Fiksel G. (2023a) Aquatic Ferrous Solutions of Prebiotic Mineral Salts as Strong UV Protectants and Possible Loci of Life Origin. Astrobiology, 23, 741–745.

Subbotin V. and Fiksel G. (2023b) Exploring the Lipid World hypothesis: A novel scenario of self-sustained Darwinian evolution of the liposomes. Astrobiology, 23, 344–357.

